# Oligomerization of the Human Adenosine A_2A_ Receptor is Driven by the Intrinsically Disordered C-terminus

**DOI:** 10.1101/2020.12.21.423144

**Authors:** Khanh D. Q. Nguyen, Michael Vigers, Eric Sefah, Susanna Seppälä, Jennifer P. Hoover, Nicole S. Schonenbach, Blake Mertz, Michelle A. O’Malley, Songi Han

## Abstract

G protein-coupled receptors (GPCRs) have long been shown to exist as oligomers with functional properties distinct from those of the monomeric counterparts, but the driving factors of GPCR oligomerization remain relatively unexplored. In this study, we focus on the human adenosine A_2A_ receptor (A_2A_R), a model GPCR that forms oligomers both *in vitro* and *in vivo*. Combining experimental and computational approaches, we discover that the intrinsically disordered C-terminus of A_2A_R drives the homo-oligomerization of the receptor. The formation of A_2A_R oligomers declines progressively and systematically with the shortening of the C-terminus. Multiple interaction sites and types are responsible for A_2A_R oligomerization, including disulfide linkages, hydrogen bonds, electrostatic interactions, and hydrophobic interactions. These interactions are enhanced by depletion interactions along the C-terminus, forming a tunable network of bonds that allow A_2A_R oligomers to adopt multiple interfaces. This study uncovers the disordered C-terminus as a prominent driving factor for the oligomerization of a GPCR, offering important guidance for structure-function studies of A_2A_R and other GPCRs.

## INTRODUCTION

G protein-coupled receptors (GPCRs) have long been studied as monomeric units, but accumulating evidence demonstrates that these receptors can also form homo- and hetero-oligomers with far-reaching functional implications. The properties emerging from these oligomers can be distinct from those of the monomeric protomers in ligand binding(1–4), G protein coupling(5–9), downstream signaling(10–13), and receptor internalization/desensitization(14–16). With the vast number of genes identified in the human genome(17), GPCRs are able to form a daunting number of combinations with unprecedented functional consequences. The existence of this intricate network of interactions among GPCRs presents major challenges and opportunities for the development of novel therapeutic approaches(18–23). Hence, it is crucial to identify the driving factors that govern the oligomerization of GPCRs, such that the properties of GPCR oligomers can be understood.

GPCR oligomers with multiple interfaces(24–28) can give rise to myriad ways by which these complexes can be formed and their functions modulated. In the crystal structure of the turkey β_1_-adrenergic receptor (β_1_AR), the receptor appears to dimerize via two different interfaces, one formed via TM4/TM5 (transmembrane domains 4/5) and the other via TM1/TM2/H8 (helix 8) contacts(29). Similarly, in the crystal structure of the antagonist-bound μ-opioid receptor (μ-OR), the protomers also dimerize via two interfaces; however, only one of them is predicted to induce a steric hindrance that prevents activation of both protomers(30), hinting at interface-specific functional consequences. A recent computational study predicted that the adenosine A_2A_ receptor (A_2A_R) forms homodimers via three different interfaces and that the resulting dimeric architectures can modulate receptor function in different or even opposite ways(27). All of the above-mentioned interfaces are symmetric, meaning that the two protomers are in face-to-face orientations, hence forming strictly dimers. Asymmetric interfaces, reported in M_3_ muscarinic receptor(31), rhodopsin(32–34), and opsin(34), are in contrast formed with the protomers positioning face-to-back, possibly enabling the association of higher-order oligomers.

Not only do GPCRs adopt multiple oligomeric interfaces, but various studies also suggest that these interfaces may dynamically rearrange to activate receptor function(35). According to a recent computational study, A_2A_R oligomers can adopt eight different interfaces that interconvert when the receptor is activated or when there are changes in the local membrane environment(24). Similarly, a recent study that combined experimental and computational data proposed that neurotensin receptor 1 (NTS_1_R) dimer is formed by “rolling” interfaces that co-exist and interconvert when the receptor is activated(36). Clearly, meaningful functional studies of GPCRs require exploring their dynamic, heterogeneous oligomeric interfaces.

The variable nature of GPCR oligomeric interfaces suggests that protomers of GPCR oligomers may be connected by tunable interactions. In this study, we explore the role of an intrinsically disordered region (IDR) of a model GPCR that could engage in diverse non-covalent interactions, such as electrostatic interactions, hydrogen bonds, or hydrophobic interactions. These non-covalent interactions are readily tunable by external factors, such as pH, salts, and solutes, and further can be entropically stabilized by depletion interactions(37–39), leading to structure formation and assembly(40–47). In a system where large protein molecules and small solute particles typically coexist in solution, assembly of the protein molecules causes their excluded volumes to overlap and the solvent volume accessible to the solutes to increase, raising the entropy of the system. The type and concentration of solutes or ions can also remove water from the hydration shell around the proteins, further enhancing entropy-driven protein-protein association in what is known as the hydrophobic effect(48). This phenomenon is applied in the precipitation of proteins upon addition of so-called salting-out ions according to the Hofmeister series(49). The ability of IDRs to readily engage in these non-covalent interactions motivates our focus on the potential role of IDRs in driving GPCR oligomerization.

The cytosolic carboxy (C-)terminus of GPCRs is usually an IDR(50, 51). Varying in length among different GPCRs, the C-terminus is commonly removed in structural studies of GPCRs to enhance receptor stability and conformational homogeneity. A striking example is A_2A_R, a model GPCR with a particularly long, 122-residue, C-terminus that is truncated in all published structural biology studies(24, 27, 52–59). However, evidence is accumulating that such truncations—shown to affect GPCR downstream signaling(60–62)—may abolish receptor oligomerization(63, 64). A study using immunofluorescence has demonstrated that C-terminally truncated A_2A_R does not show protein aggregation or clustering on the cell surface, a process readily observed in the wild-type form(65). Our recent study employing a tandem three-step chromatography approach uncovered the impact of a single residue substitution of a C-terminal cysteine, C394S, in reducing the receptor homo-oligomerization *in vitro*(63). In the context of heteromerization, mass spectrometry and pull-down experiments have demonstrated that A_2A_R-D_2_R dimerization occurs via direct electrostatic interactions between the C-terminus of A_2A_R and the third intracellular loop of D_2_R(66). These results all suggest that the C-terminus may participate in A_2A_R oligomer formation. However, no studies to date have directly and systematically investigated the role of the C-terminus, or any IDRs, in GPCR oligomerization.

This study focuses on the homooligomerization of the human adenosine A_2A_R, a model GPCR, and seeks to address: (i) whether the C-terminus engages in A_2A_R oligomerization, and if so, (ii) whether the C-terminus forms multiple oligomeric interfaces. We use size-exclusion chromatography (SEC) to assess the oligomerization levels of A_2A_R variants with strategic C-terminal modifications: mutations of a cysteine residue C394 and a cluster of charged residues ^355^ERR^357^, as well as systematic truncations at eight different sites along its length. We complemented our experimental study with an independent molecular dynamics study of A_2A_R dimers of five C-terminally truncated A_2A_R variants designed to mirror the experimental constructs. We furthermore examined the oligomerization level of select C-terminally modified A_2A_R variants under conditions of ionic strength ranging from 0.15 to 0.95 M. To test whether the C-termini directly and independently promote A_2A_R oligomerization, we recombinantly expressed the entire A_2A_R C-terminal segment sans the transmembrane portion of the receptor and investigated its solubility and assembly properties with increasing ion concentration and temperature. This is the first study designed to uncover the role of the intrinsically disordered C-terminus on the oligomerization of a GPCR.

## RESULTS

This study systematically investigates the role of the C-terminus on A_2A_R oligomerization and the nature of the interactions involved through strategic mutations and truncations at the C-terminus as well as modulation of the ionic strength of solvent. The experimental assessment of A_2A_R oligomerization relies on size-exclusion chromatography (SEC) analysis.

### Size Exclusion Chromatography Quantifies A_2A_R Oligomerization

We performed SEC analysis on a mixture of ligand-active A_2A_R purified from a custom synthesized antagonist affinity column (**Fig. S1A**). Distinct oligomeric species were separated and eluted in the following order: high-molecular-weight (HMW) oligomer, dimer, and monomer (**Fig. 1** and **Fig. S1B**). The population of each oligomeric species was quantified as the integral of each Gaussian from a multiple-Gaussian curve fit of the SEC signal. The reported standard errors were calculated from the variance of the fit that do not correspond to experimental errors (see **Table S1** and **Fig. S2** for SEC data corresponding to all A_2A_R variants in this study). As this study sought to identify the factors that promote A_2A_R oligomerization, the populations with oligomeric interfaces (*i.e.*, dimer and HMW oligomer) were compared with those without such interfaces (*i.e.,* monomer). Hence, the populations of the HMW oligomer and dimer were expressed relative to the monomer population in arbitrary units as monomer-equivalent concentration ratios, henceforth referred to as population levels (**Fig. 1)**.

**Figure 1.**
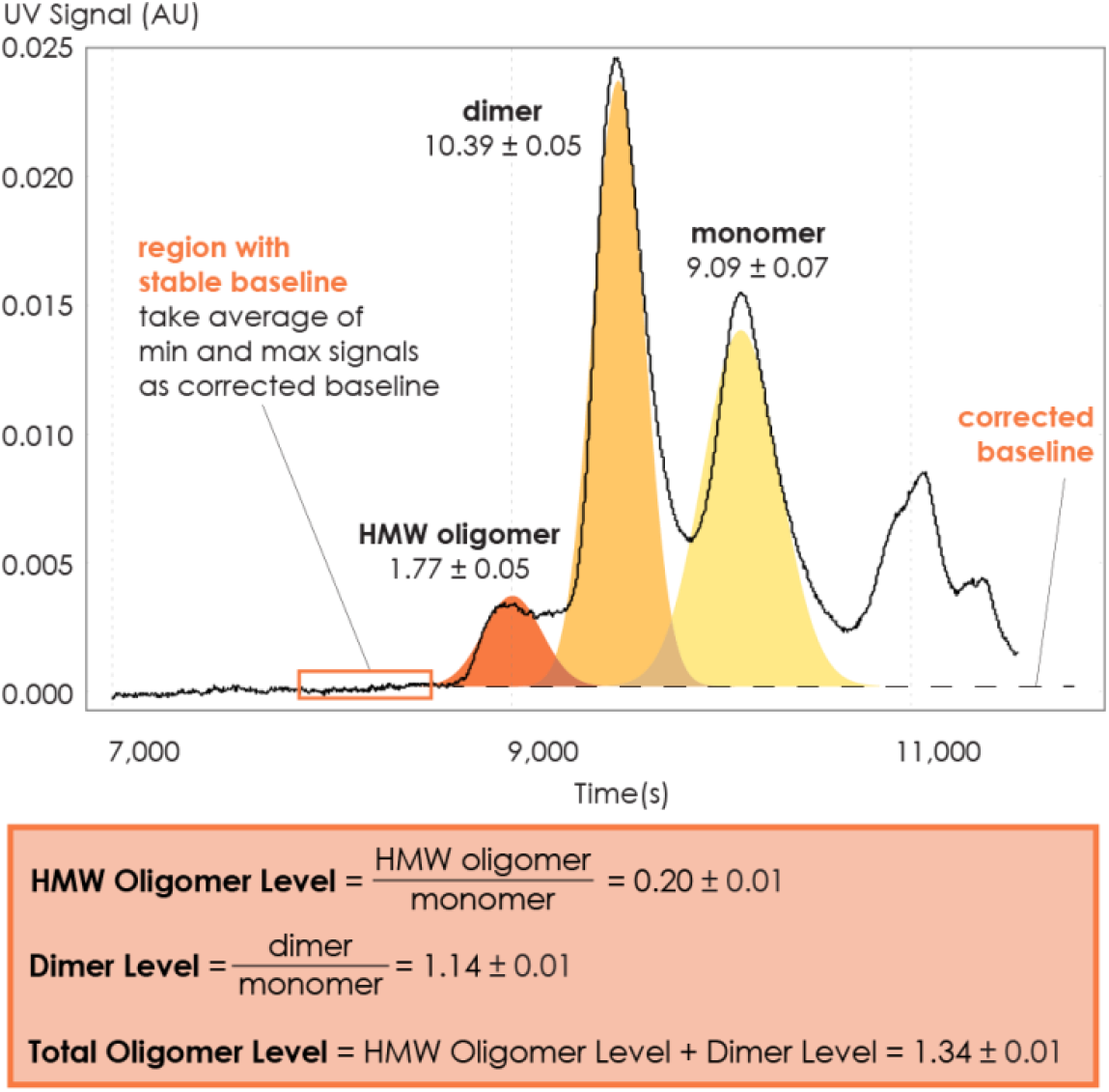
Method for collecting SEC data and assessing A_2A_R oligomerization. The SEC data is recorded every second as absorbance at 280 nm. The baseline is corrected to ensure uniform fitting and integration across the peaks. The areas under the curve, resulting from a multiple-Gaussian curve fit, express the population of each oligomeric species. The reported standard errors of integration are within a 95% confidence interval and are calculated from the variance of the fit, not experimental errors. The levels of HMW oligomer and dimer are expressed relative to the monomeric population in arbitrary units. A representative calculation defining the oligomer levels is given in the box.

### C-Terminal Amino Acid Residue C394 Contributes to A_2A_R Oligomerization

To investigate whether the C-terminus of A_2A_R is involved in receptor oligomerization, we first examined the role of residue C394, as a previous study demonstrated that the mutation C394S dramatically reduced A_2A_R oligomer levels(63). The C394S mutation was replicated in our experiments, alongside other amino acid substitutions, namely alanine, leucine, methionine or valine, generating five A_2A_R-C394X variants. The HMW oligomer and dimer levels of A_2A_R wild-type (WT) were compared with those of the A_2A_R-C394X variants. We found that the dimer level of A_2A_R-WT was significantly higher than that of the A_2A_R-C394X variants (WT: 1.14; C394X: 0.24–0.57; **Fig. 2A**). A similar result, though less pronounced, was observed when the HMW oligomer and dimer levels were considered together (WT: 1.34; C394X: 0.59–1.21; **Fig. 2A**). This suggests that residue C394 plays a role in A_2A_R oligomerization and more so in A_2A_R dimers.

**Figure 2.**
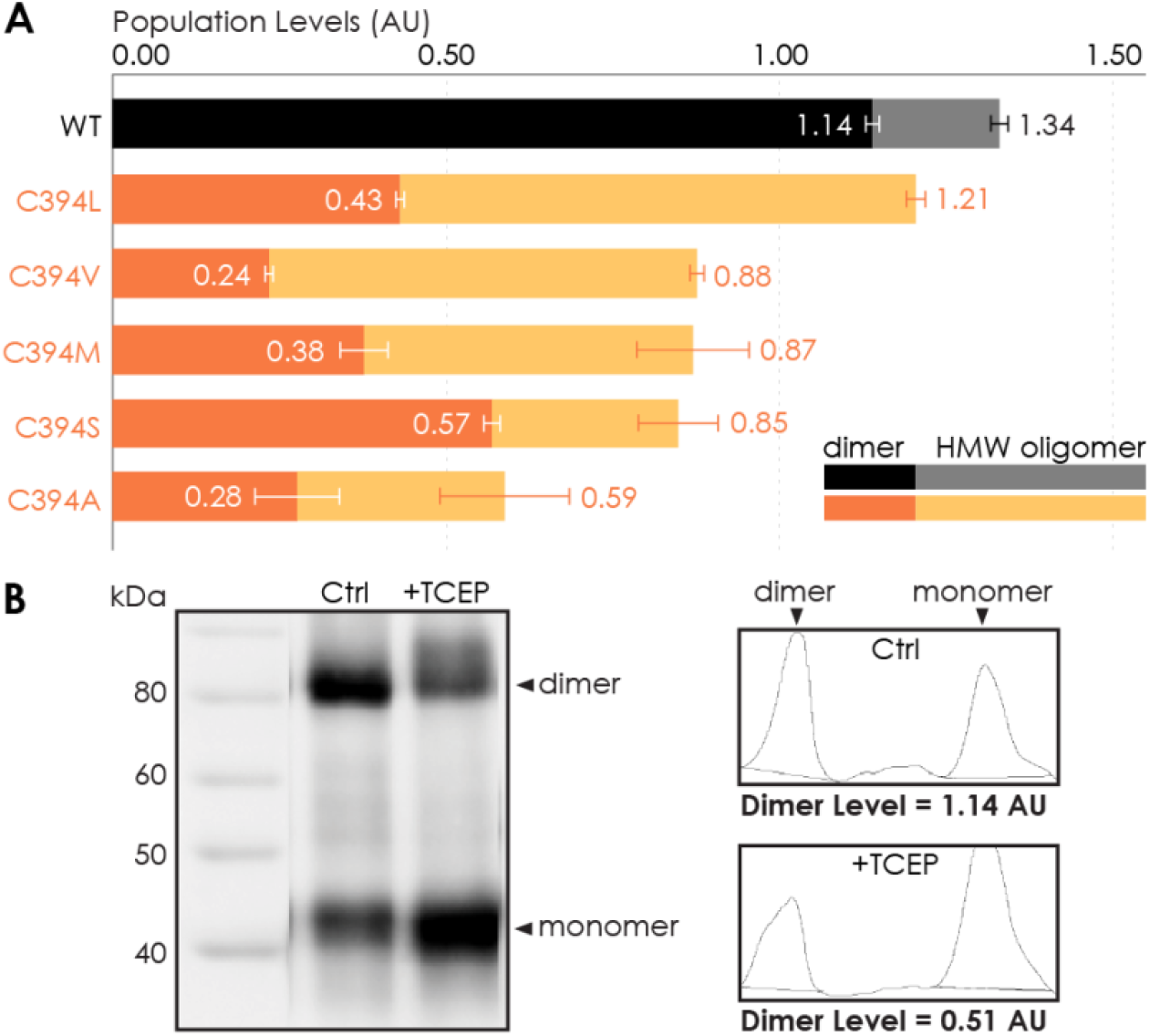
Residue C394 helps stabilize A_2A_R oligomerization via disulfide bonds. (A) The effect of C394X substitutions on A_2A_R oligomerization. The levels of dimer (dark colors) and HMW oligomer (light colors) are expressed relative to the monomeric population in arbitrary units, with reported errors calculated from the variance of the fit, not experimental variation. (B) Line densitometry of Western Blot bands on SEC-separated dimeric population with and without 5 mM TCEP. The level of dimer is expressed relative to the monomeric population in arbitrary units similarly to the SEC analysis. MagicMark protein ladder (LC5602) is used as the molecular weight standard.

To test whether residue C394 stabilizes A_2A_R dimerization by forming disulfide linkages, we incubated SEC-separated A_2A_R dimer with 5 mM of the reducing agent TCEP, followed by SDS-PAGE and Western Blotting. The population of each species was determined as the area under the densitometric trace. The dimer level was then expressed as monomer-equivalent concentration ratios in a manner similar to that of the SEC experiment described above. Upon incubation with TCEP, the dimer level of the sample decreased from 1.14 to 0.51 (**Fig. 2B**). This indicates that disulfide bond formation via residue C394 is one possible mechanism for A_2A_R dimerization. However, a significant population of A_2A_R dimer remained resistant to TCEP and C394X mutations (**Fig. 2**), suggesting that disulfide linkages are not the only driving factor of A_2A_R oligomer formation. This finding agrees with a previous study showing that residue C394 in A_2A_R dimer is still available for nitroxide spin labeling,(63) suggesting that additional interfacial sites help drive A_2A_R dimer/oligomerization.

### C-Terminus Truncation Systematically Reduces A_2A_R Oligomerization

To determine which interfacial sites in the C-terminus other than C394 drive A_2A_R dimer/oligomerization, we carried out systematic truncations at eight sites along the C-terminus (A316, V334, G344, G349, P354, N359, Q372, and P395), generating eight A_2A_R-ΔC variants (**Fig. 3A**). The A_2A_R-A316ΔC variant corresponds to the removal of the entire disordered C-terminal region as previously performed in all published structural studies(24, 27, 52–59). Using the SEC analysis described earlier (**Fig. 1**) we evaluated the HMW oligomer and dimer levels of the A_2A_R-ΔC variants relative to that of the A_2A_R full-length-wild-type (FL-WT) control. Both the dimer and the total oligomer levels of A_2A_R decreased progressively with the shortening of the C-terminus, with almost no oligomerization detected upon complete truncation of the C-terminus at site A316 (**Fig. 3B**). This result shows that the C-terminus drives A_2A_R oligomerization, with multiple potential interaction sites positioned along much of its length.

**Figure 3.**
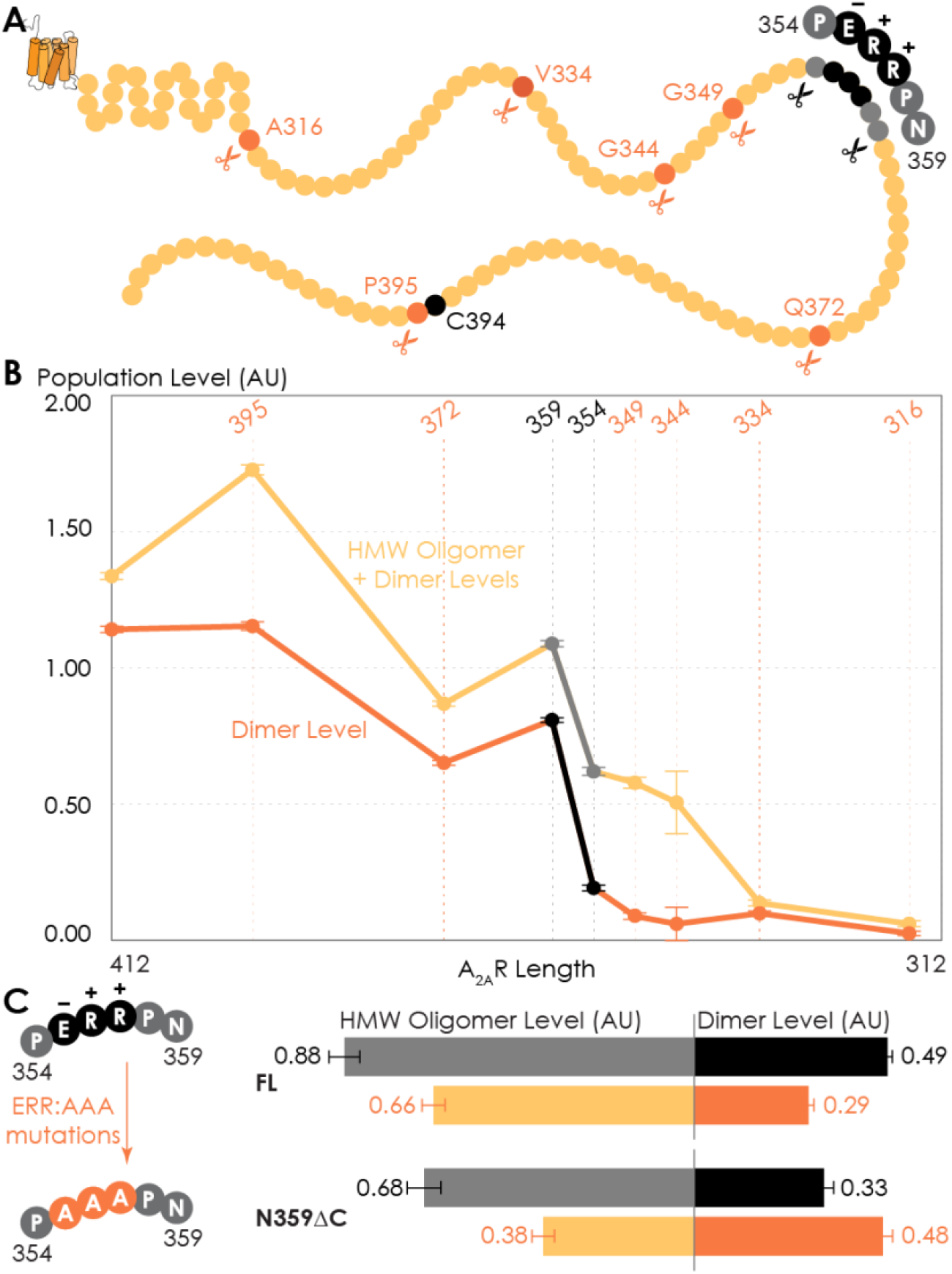
Truncating the C-terminus systematically affects A_2A_R oligomerization. (A) Depiction of where the truncation points are located on the C-terminus, with region 354–359 highlighted (in black) showing critical residues. (B) The levels of dimer and HMW oligomer are expressed relative to the monomeric population as an arbitrary unit and plotted against the residue number of the truncation sites, with reported errors calculated from the variance of the fit, not experimental variation. Region 354–359 is emphasized (in black and gray) due to a drastic change in the dimer and HMW oligomer levels. (C) The dependence of A_2A_R oligomerization on three consecutive charged residues ^355^ERR^357^. The substitution of residues ^355^ERR^357^ to 355AAA357 is referred to as the ERR:AAA mutations. The levels of dimer and HMW oligomer are expressed relative to the monomeric population as an arbitrary unit, with reported errors calculated from the variance of the fit, not experimental variation.

Interestingly, there occurred a dramatic decrease in the dimer level between the N359 and P354 truncation sites, from a value of 0.81 to 0.19, respectively (**Fig. 3B**). A similar result, though less pronounced, was observed on the total oligomer level, with a decrease from 1.09 to 0.62 for the N359 and P354 truncation sites, respectively (**Fig. 3B**). Clearly, the C-terminal segment encompassing residues 354–359 (highlighted in black in **Fig. 3A**) is a key constituent of the A_2A_R oligomeric interface.

Since segment 354–359 contains three consecutive charged residues (^355^ERR^357^; **Fig. 3A**), which could be involved in electrostatic interactions, we hypothesized that this ^355^ERR^357^ cluster could strengthen inter-protomer A_2A_R-A_2A_R association. To test this hypothesis, residues ^355^ERR^357^ were substituted by 355AAA357 on A_2A_R-FL-WT and A_2A_R-N359ΔC to generate A_2A_R-ERR:AAA variants (**Fig. 3C**). We then compared the HMW oligomer and dimer levels of the resulting variants with controls (same A_2A_R variants but without the ERR:AAA mutations). We found that the ERR:AAA mutations had varied effects on the dimer level: decreasing for A_2A_R-FL-WT (ctrl: 0.49; ERR:AAA: 0.29) but increasing for A_2A_R-N359ΔC (ctrl: 0.33; ERR:AAA: 0.48) (**Fig. 3C**). In contrast, the ERR:AAA mutations reduced the HMW oligomer level of both A_2A_R-FL-WT (ctrl: 0.88; ERR:AAA: 0.66) and A_2A_R-N359ΔC (ctrl: 0.68; ERR:AAA: 0.38) (**Fig. 3C**). Consistently, the ERR:AAA mutation lowered the total oligomer level of both A_2A_R-FL-WT (ctrl: 1.37; ERR:AAA: 0.94) and A_2A_R-N359ΔC (ctrl: 1.01; ERR:AAA: 0.85) (**Fig. 3C**). These results suggest that the charged residues ^355^ERR^357^ participate in A_2A_R oligomerization, with a greater effect in the context of a longer C-terminus and for higher-order oligomer formation. The question then arises as to what types of interactions are formed along the C-terminus that help stabilize A_2A_R oligomerization.

### C-Terminus Truncation Disrupts Complex Network of Non-Bonded Interactions Necessary for A_2A_R Dimerization

Given that the structure of A_2A_R dimers or oligomers are unknown, we next used molecular dynamics (MD) simulations to seek molecular-level insights into the role of the C-terminus in driving A_2A_R dimerization and to determine the specific interaction types and sites involved in this process. First, to explore A_2A_R dimeric interface, we performed coarse-grained (CG) MD simulations, which can access the length and time scales relevant to membrane protein oligomerization, albeit at the expense of atomic-level details. We carried out a series of CGMD simulations on five A_2A_R-ΔC variants designed to mirror the experiments by systematic truncation at five sites along the C-terminus (A316, V334, P354, N359, and C394). Our results revealed that A_2A_R dimers were formed with multiple interfaces, all involving the C-terminus (**Fig. 4A** and **S3A**).

**Figure 4.**
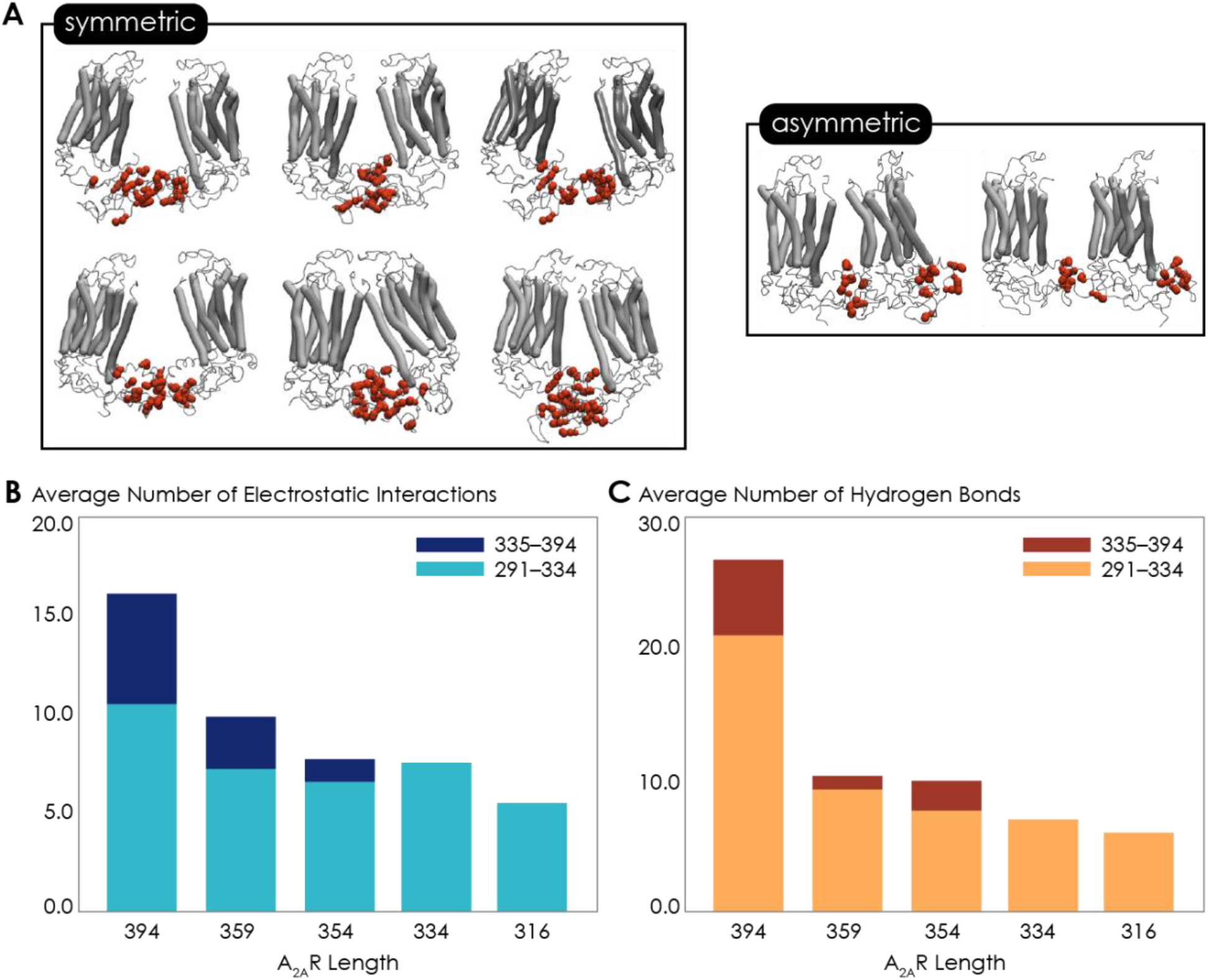
Non-bonded interactions of the extended C-terminus of A_2A_R play a critical role in stabilization of the dimeric interface. **(A)** Dimer configurations from cluster analysis in GROMACS of the 394-residue variant identify two major clusters involving either 1) the C-terminus of one protomer and the C-terminus, ICL2, and ICL3 of the second protomer or 2) the C-terminus of one protomer and ICL2, ICL3, and ECL2 of the second protomer. Spheres: residues forming intermolecular electrostatic contacts. **(B)** Average number of residues that form electrostatic contacts as a function of sequence length of A_2A_R. **(C)** Average number of residues that form hydrogen bonds as a function of sequence length of A_2A_R.

The vast majority of A_2A_R dimers were symmetric, with the C-termini of the protomers directly interacting with each other. A smaller fraction of the dimers had asymmetric orientations, with the C-terminus of one protomer interacting with other parts of the other protomer, such as ICL2 (the second intracellular loop), ICL3, and ECL2 (the second extracellular loop) (**Fig. 4A**).

Our observation of multiple A_2A_R oligomeric interfaces, consistent with previous studies(24, 27), suggests that tunable, non-covalent intermolecular interactions are involved in receptor dimerization. We dissected two key non-covalent interaction types: electrostatic and hydrogen bonding interactions. (The criteria for designating inter-A_2A_R contacts as electrostatic interactions or hydrogen bonds are described in detail in **Materials and Methods.**) Electrostatic interactions were calculated from CGMD simulations. Hydrogen bonds were quantified from atomistic MD simulation, given that the CG model merges all hydrogens into a coarse-grained bead and hence cannot report on hydrogen bonds. This analysis was performed on the symmetric dimers as they constituted the majority of the assemblies. With the least truncated A_2A_R variant containing the longest C-terminus, A_2A_R-C394ΔC, we observed an average of 15.9 electrostatic contacts (**Fig. 4B**) and 26.7 hydrogen bonds (**Fig. 4C**) between the C-termini of the protomers. This result shows that both electrostatic interactions and hydrogen bonds play important roles in A_2A_R dimer formation.

Upon further C-terminus truncation, the average number of both electrostatic contacts and hydrogen bonds involving C-terminal residues progressively declined, respectively reaching 5.4 and 6.0 for A_2A_R-A316ΔC (in which the disordered region of the C-terminus is removed) (**Fig. 4B** and **4C**). This result is consistent with the experimental result, which demonstrated a progressive decrease of A_2A_R oligomerization with the shortening of the C-terminus (**Fig. 3B**). Interestingly, upon systematic truncation of the C-terminal segment 335–394, we observed in segment 291–334 a steady decrease in the average number of electrostatic contacts, from 10.4 to 7.4 (**Fig. 4B**). This trend was even more pronounced with hydrogen bonding contacts involving segment 291–334 decreasing drastically from 21.0 to 7.0 as segment 335–394 was gradually removed (**Fig. 4C**). This observation, namely that truncation of a C-terminal segment reduces inter-A_2A_R contacts elsewhere along the C-terminus, indicates that a cooperative mechanism of dimerization exists, in which an extended C-terminus of A_2A_R stabilizes inter-A_2A_R interactions near the heptahelical bundles of the dimeric complex. Besides the intermolecular interactions, we also identified a network of intramolecular salt bridges involving residues on the C-termini, including cluster ^355^ERR^357^ (**Fig. 7A**). These results demonstrate that A_2A_R dimers can be formed via multiple interfaces predominantly in symmetric orientations, facilitated a cooperative network of electrostatic interactions and hydrogen bonds along much of its C-terminus.

### Ionic Strength Modulates Oligomerization of C-Terminally Truncated A_2A_R Variants

So far, we have demonstrated that the C-terminus clearly plays a role in forming A_2A_R oligomeric interfaces. However, the driving factors of A_2A_R oligomerization remain unknown. The variable nature of A_2A_R oligomeric interfaces suggests that the main driving forces must be non-covalent interactions, such as electrostatic interactions and hydrogen bonds as identified by the above MD simulations. Modulating the solvent ionic strength is an effective method to identify the types of non-covalent interaction(s) at play. Specifically, with increasing ionic strength, electrostatic interactions can be weakened (based on Debye-Hückel theory, most electrostatic bonds at a distance greater than 5 Å are screened out at an ionic strength of 0.34 M at 4°C), depletion interactions are enhanced with salting-out salts, and hydrogen bonds remain relatively impervious. For this reason, we subjected various A_2A_R variants (FL-WT, FL-ERR:AAA, N359ΔC, and V334ΔC) to ionic strength ranging from 0.15 to 0.95 M by adding NaCl (buffer composition shown in **Table 1)**. The HMW oligomer and dimer levels of the four A_2A_R variants were determined and plotted as a function of ionic strengths.

**Table 1.** Calculations regarding composition of the buffers used in the experiments where salt concentrations are varied. Only NaCl concentration (in bold) is varied to achieve the different ionic strengths.

| *Buffers* | *Components* | *Conc.*<br>*(mM)* | *Ionic Strength*<br>*(mM)* |
| --- | --- | --- | --- |
| *0.15 M Ionic Strength* | **NaCl** | **0** | **0** |
|  | NaH<sub>2</sub>PO<sub>4</sub> | 4 | 4 |
|  | Na<sub>2</sub>HPO<sub>4</sub> | 49 | 146 |
| *0.45 M Ionic Strength* | **NaCl** | **300** | **300** |
|  | NaH<sub>2</sub>PO<sub>4</sub> | 4 | 4 |
|  | Na<sub>2</sub>HPO<sub>4</sub> | 49 | 146 |
| *0.95 M Ionic Strength* | **NaCl** | **800** | **800** |
|  | NaH<sub>2</sub>PO<sub>4</sub> | 4 | 4 |
|  | Na<sub>2</sub>HPO<sub>4</sub> | 49 | 146 |

The low ionic strength of 0.15 M should not affect hydrogen bonds or electrostatic interactions, if present. We found that the dimer and total oligomer levels of all four variants were near zero (**Fig. 5**). This is a striking observation, as it already excludes electrostatic and hydrogen-bonding interactions as the dominant force for A_2A_R association. The question remains whether depletion interactions could be involved.

**Figure 5.**
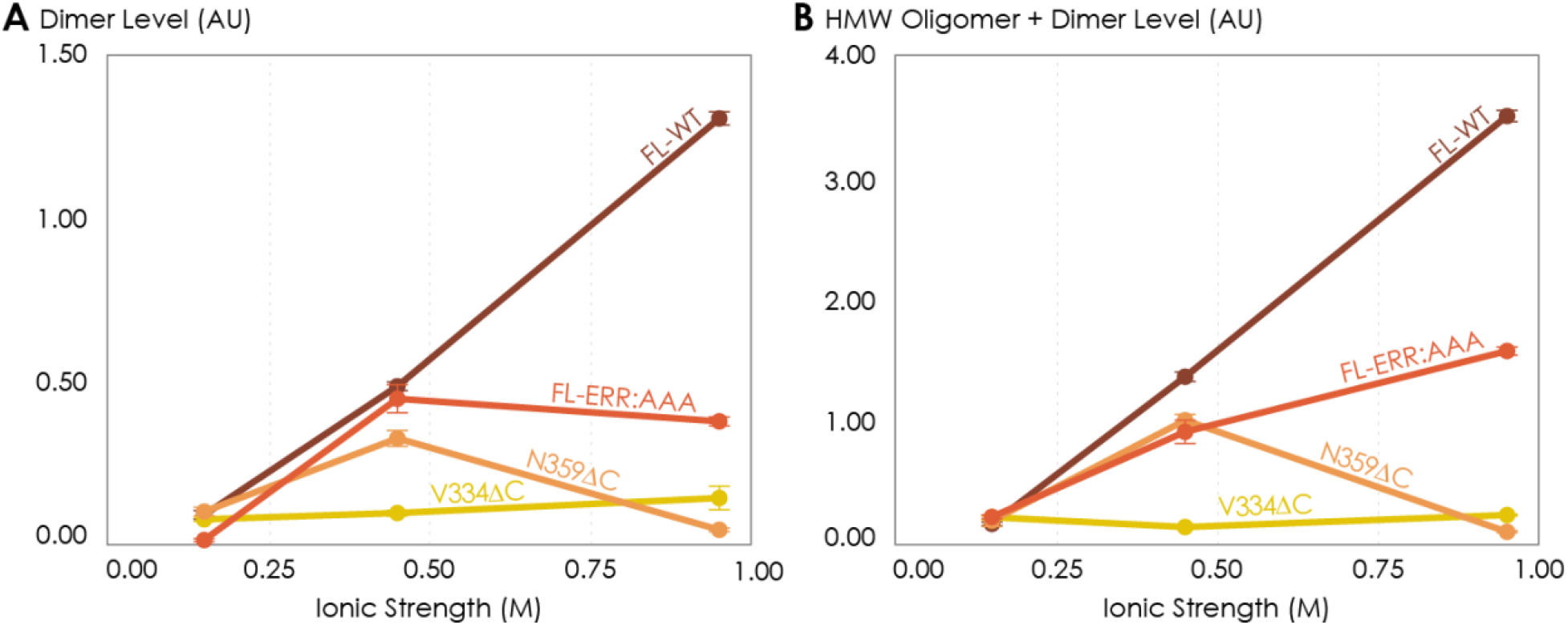
The effects of ionic strength on the oligomerization of various A_2A_R variants reveal the involvement of depletion interactions. The levels of dimer and HMW oligomer are expressed relative to the monomeric population as an arbitrary unit and plotted against ionic strength, with reported errors calculated from the variance of the fit, not experimental variation. NaCl concentration is varied to achieve ionic strengths of 0.15, 0.45, and 0.95 M.

At higher ionic strengths of 0.45 M and 0.95 M, the dimer and total oligomer levels of A_2A_R-V334ΔC still remained near zero (**Fig. 5)**. In contrast, we observed a progressive and significant increase in the dimer and total oligomer levels of A_2A_R-FL-WT with increasing ionic strength (**Fig. 5)**. This result indicates A_2A_R oligomerization must be driven by depletion interactions, which are enhanced with increasing ionic strength, and that these interactions involve the C-terminal segment after residue V334.

Upon closer examination, we recognize that at the very high ionic strength of 0.95 M, the increase in the dimer and total oligomer levels was robust for A_2A_R-FL-WT, but less pronounced for A_2A_R-FL-ERR:AAA (**Fig. 5**). Furthermore, this high ionic strength even had an opposite effect on A_2A_R-N359ΔC, with both its dimer and total oligomer levels abolished (**Fig. 5**). These results indicate that the charged cluster ^355^ERR^357^ and the C-terminal segment after residue N359 are required for depletion interactions to promote A_2A_R oligomerization to the full extent.

Taken together, we demonstrated that A_2A_R oligomerization is more robust when the C-terminus is fully present and the ionic strength is higher, suggesting that depletion interactions via the C-terminus are a strong driving factor of A_2A_R oligomerization. The question then arises whether such depletion interactions are the result of the C-termini directly interacting with one another, necessitating an experiment that investigates the behavior of A_2A_R C-terminus sans the transmembrane domains.

### The Isolated A_2A_R C-Terminus Is Prone to Aggregation

To test whether A_2A_R oligomerization is driven by direct depletion interactions among the C-termini of the protomers, we assayed the solubility and assembly properties of the stand-alone A_2A_R C-terminus—an intrinsically disordered peptide—sans the upstream transmembrane regions. Since depletion interactions can be manifested via the hydrophobic effect(48), we examined whether this effect can cause A_2A_R C-terminal peptides to associate.

It is an active debate(67) whether the hydrophobic effect can be promoted or suppressed by ions with salting-out or salting-in tendency, respectively(68–70). We increased the solvent ionic strength using either sodium (salting-out) or guanidinium (salting-in) ions and assessed the aggregation propensity of the C-terminal peptides using UV-Vis absorption at 450 nm. We first observed the behavior of the C-terminus with increasing salting-out NaCl concentrations. At NaCl concentrations below 1 M, the peptide was dominantly monomeric, despite showing slight aggregation at NaCl concentrations between 250–500 mM (**Fig. 6A**). At NaCl concentrations above 1 M, A_2A_R C-terminal peptides strongly associated into insoluble aggregates (**Fig. 6A**). Consistent with the observations made with the intact receptor (**Fig. 5**), A_2A_R C-terminus showed the tendency to progressively precipitate with increasing ionic strengths, suggesting that depletion interactions drive the association and precipitation of the peptides. We next observed the behavior of the C-terminus with increasing concentrations of guanidine hydrochloride (GdnHCl), which contains salting-in cations that do not cause proteins to precipitate and instead facilitate the solubilization of proteins(71, 72). Our results demonstrated that the A_2A_R C-terminus incubated in 4 M GdnHCl showed no aggregation propensity (**Fig. 6A)**, validating our expectation that depletion interactions are not enhanced by salting-out salts. These observations demonstrate that the C-terminal peptide in and of itself can directly interact with other C-terminal peptides to form self-aggregates in the presence of ions, and presumably solutes, that have salting-out effects.

**Figure 6.**
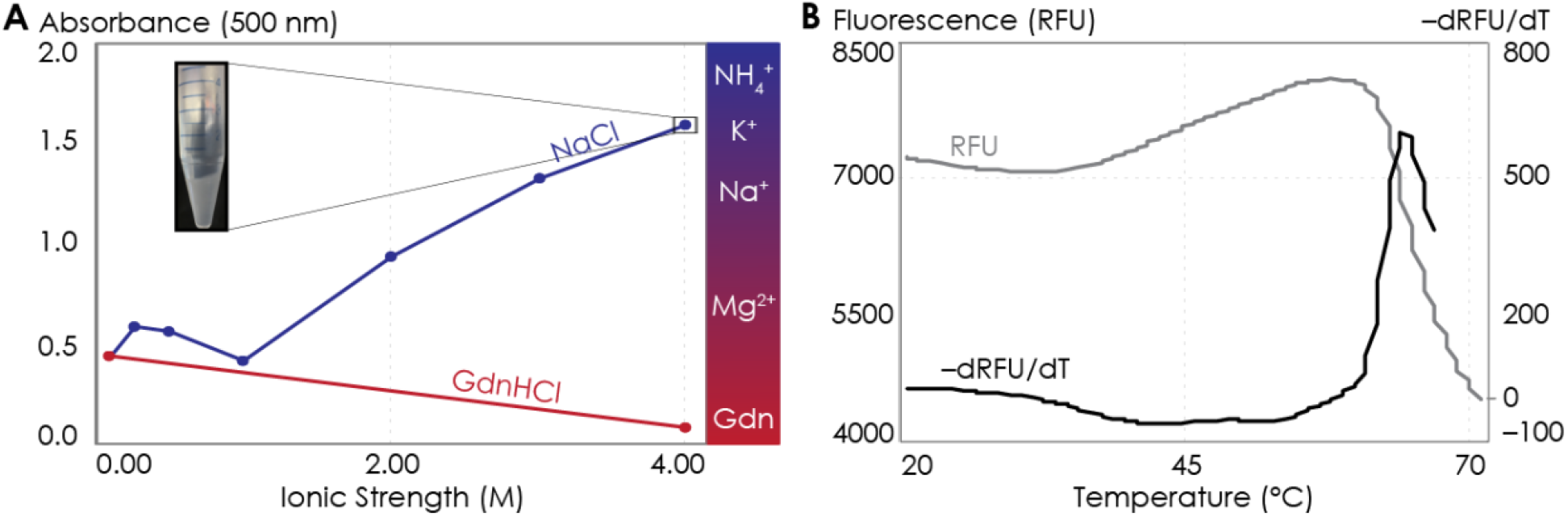
The A_2A_R C-terminus is prone to aggregation. (A) Absorbance at 500 nm of the A_2A_R C-terminus in solution, with NaCl and GdnHCl concentrations varied to achieve ionic strengths 0–4 M. Inset: the solution at ionic strength 4 M achieved with NaCl. The Hofmeister series is provided to show the ability of cations to salt out (blue) or salt in (red) proteins. (B) SYPRO orange fluorescence of solutions containing the A_2A_R C-terminus as the temperature was varied from 20 to 70°C (grey). The change in fluorescence, measured in relative fluorescence unit (RFU), was calculated by taking the first derivative of the fluorescence curve (black).

**Figure 7.**
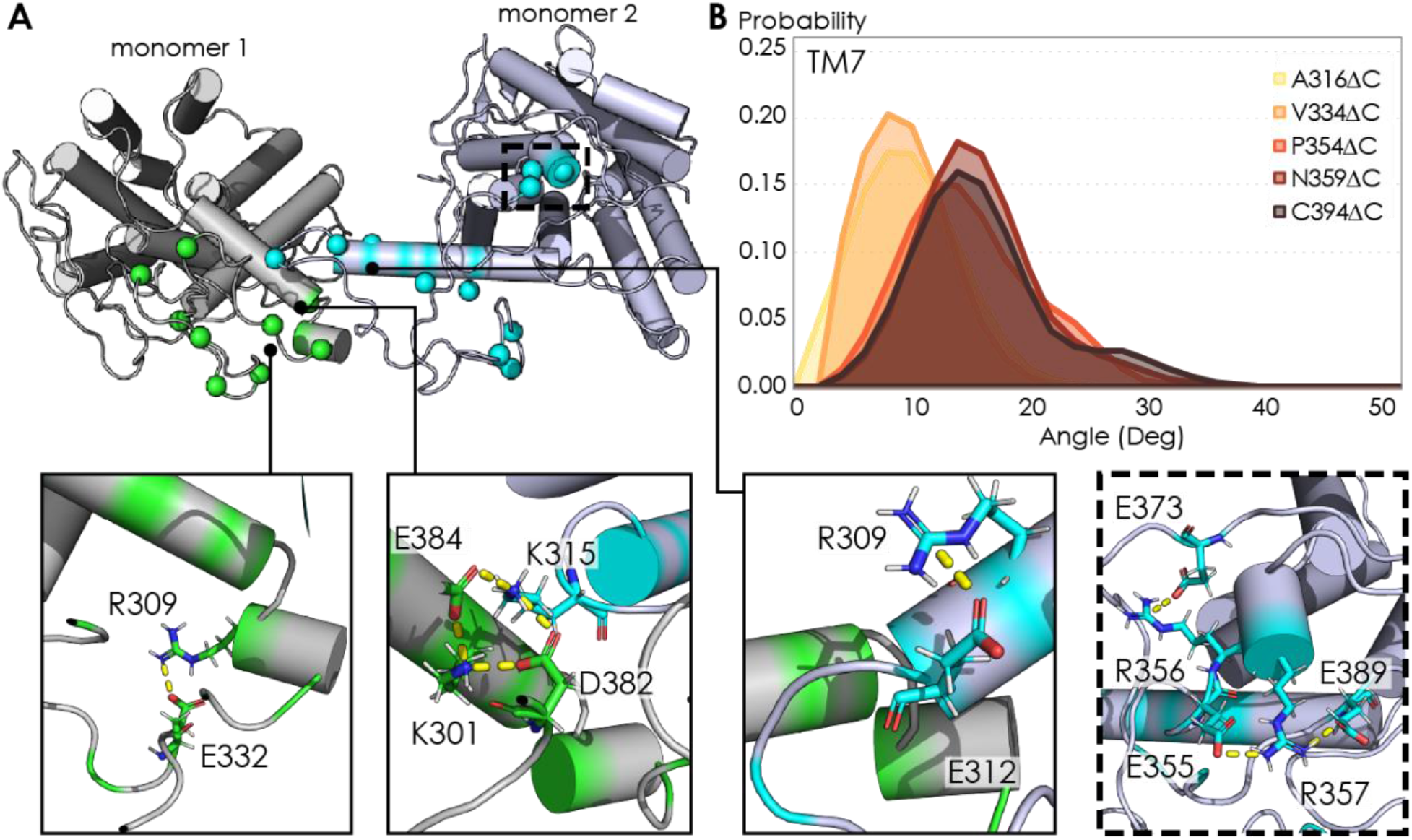
**(A)** Representative snapshot of A_2A_R-C394ΔC dimers shows salt bridge formation between a sample trajectory. The insets are close-ups of the salt bridges, which can be both intra- and intermolecular. The last inset shows a network of salt bridges with the charged cluster ^355^ERR^357^ involved. **(B)** Helical tilt angles for TM7 helix in A_2A_R as a function of protein length. Systematic truncations of the C-terminus lead to rearrangement of the heptahelical bundle. The participation of the C-terminus in A_2A_R dimerization increases the tilting of the TM7 domain, which is in closest proximity to the C-terminus.

Attractive hydrophobic interactions among the hydrophobic residues are further enhanced by water solvating the protein having more favorable interactions with other water molecules, ions or solutes than with the protein, here the truncated C-terminus(73–75). We explored the possible contribution of hydrophobic interactions to the aggregation of the C-terminal peptides using differential scanning fluorimetry (DSF). In particular, we gradually increased the temperature to melt the C-terminal peptides, exposing any previously buried hydrophobic residues (**Fig. S4A**) which then bound to the SYPRO orange fluorophore, resulting in an increase in fluorescence signal. Our results showed that as the temperature increased, a steady rise in fluorescence was observed (**Fig. 6B**), indicating that multiple hydrophobic residues were gradually exposed to the SYPRO dye. However, at approximately 65°C, the melt peak signal was abruptly quenched (**Fig. 6B**), indicating that the hydrophobic residues were no longer exposed to the dye. This observation suggests that, at 65°C, enough hydrophobic residues in the C-terminal peptides were exposed such that they collapsed on one another (thus expelling the bound dye molecules), resulting in aggregation. Clearly, the hydrophobic effect can cause A_2A_R C-terminal peptides to directly associate. These results demonstrate that A_2A_R oligomer formation can be driven by depletion interactions among the C-termini of the protomers.

## DISCUSSION

The key finding of this study is that the C-terminus of A_2A_R, removed in all previously published structural studies of this receptor, is directly responsible for receptor oligomerization. Using a combination of experimental and computational approaches, we demonstrate that the C-terminus drives A_2A_R oligomerization via a combination of disulfide linkages, hydrogen bonds, electrostatic interactions, and hydrophobic interactions. This diverse combination of interactions is greatly enhanced by depletion interactions, forming a network of malleable bonds that give rise to the existence of multiple A_2A_R oligomeric interfaces.

The intermolecular disulfide linkages associated with residue C394 play a role in A_2A_R oligomerization. However, it is unclear which cysteine on the second protomer is linked to this cysteine. A previous study showed that residue C394 in A_2A_R dimer is available for nitroxide spin labeling(63), suggesting that some of these disulfide bonds may be between residue C394 and another cysteine in the hydrophobic core of A_2A_R that do not form intramolecular disulfide bonds(76–78). Many examples exist where disulfide linkages help drive GPCR oligomerization, including the CaR-mGluR_1_ heterodimer(79), homodimers of mGluR_5_(80), M_3_R(81), V_2_R(82), 5-HT_4_R(83) and 5-HT_1D_R(84), and even higher-order oligomers of D_2_R(85). However, although unconventional cytoplasmic disulfide bonds have been reported(86, 87), no study has shown how such linkages would be formed *in vivo*, as the cytoplasm lacks the conditions and machinery required for disulfide bond formation(88–91). Nevertheless, residue C394 is highly conserved and a C-terminal cysteine is almost always present among A_2A_R homologs(92), suggesting that this cysteine cannot be excluded for serving an important role *in vivo*.

The electrostatic interactions that stabilize A_2A_R oligomer formation come from multiple sites along the C-terminus. From a representative snapshot of a A_2A_R-C394ΔC dimer from our MD simulations (**Fig. 7A**), we could visualize not only the intermolecular interactions calculated from the CGMD simulations (**Fig. 4B**), but also intramolecular salt bridges. In particular, the ^355^ERR^357^ cluster of charged residues lies distal from the dimeric interface, yet still forms several salt bridges (**Fig. 7A**, inset). This observation is supported by our experimental results showing that substituting this charged cluster with alanines reduces the total A_2A_R oligomer levels (**Fig. 3C**). However, it is unclear how such salt bridges involving this ^355^ERR^357^ cluster are enhanced by depletion interactions (**Fig. 5**), as electrostatic interactions are usually screened out at high ionic strengths. In our MD simulations, we also observed networks of salt bridges along the dimeric interface, for example between K315 of one monomer and D382 and E384 of the other monomer (**Fig. 7A**, inset). The innate flexibility of the C-terminus could facilitate the formation of such salt bridges, which then acts as a potential scaffold to stabilize A_2A_R dimers.

We also found that depletion interactions can enhance the diversity of interactions that stabilize A_2A_R oligomer formation (**Fig. 5** and **6**). Depletion interactions could be the key factor to the cooperative mechanism by which A_2A_R oligomerization occurs. As revealed by our MD simulations, an increasing number of contacts are formed along segment 291–334 when the rest of C-terminus is present (**Fig. 4B** and **4C)**. As more of the C-terminus is preserved, the greater extent of depletion interactions limits the available dimer arrangements, forcing segment 291–334 into an orientation that optimizes intermolecular interactions.

Our finding that A_2A_R forms homo-oligomers via multiple interfaces (**Fig. 4A**) agrees with the increasing number of studies reporting multiple and interconverting oligomeric interfaces in A_2A_R and other GPCRs(24–36). When translated to *in vivo* situations, GPCR oligomers can also transiently associate and dissociate(93–96). Such fast conformational changes require that the oligomeric interfaces be formed by interactions that can easily be modulated. This is consistent with our study, which demonstrates that depletion interactions via the intrinsically disordered, malleable C-terminus drive A_2A_R oligomerization. Because depletion interactions can be readily tuned by environmental factors, such as ionic strength, molecular crowding, and temperature, the formation of GPCR oligomeric complexes could be dynamically modulated in response to environmental cues to regulate receptor function.

Not only did we find multiple A_2A_R oligomeric interfaces, we also found that these interfaces can be either symmetric or asymmetric. This finding is supported by a growing body of evidence that there exists both symmetric and asymmetric oligomeric interfaces for A_2A_R(24) and many other GPCRs. Studies using various biochemical and biophysical techniques have shown that heterotetrameric GPCR complexes can be formed by dimers of dimers, including μOR-δOR(97), CXC_4_R-CC_2_R(98), CB_1_R/D_2_R(99) as well as those involving A_2A_R, such as A_1_R-A_2A_R(61, 100) and A_2A_R-D_2_R(101). The quaternary structures identified in these studies required specific orientations of each protomer, with the most viable model involving a stagger of homodimers with symmetric interfaces(102). On the other hand, since symmetric interfaces limit the degree of receptor association to dimers, the HMW oligomer of A_2A_R observed in this(24) and other studies(63, 103) can only be formed via asymmetric interfaces. It is indeed tempting to suggest that the formation of the HMW oligomer of A_2A_R may even arise from combinations of different interfaces. In any case, the wide variation of GPCR oligomerization requires the existence of both symmetric and asymmetric oligomeric interfaces.

In the case of A_2A_R, displacement of the transmembrane domains have been demonstrated to be the hallmark of receptor activation(104–107). However, no studies have linked receptor oligomerization with the arrangement of the TM bundles in A_2A_R. Our MD simulations revealed that C-terminus truncation resulted in structural changes in the heptahelical bundles of A_2A_R dimers. Specifically, as more of the C-terminus was preserved, we observed a progressive increase in the helical tilt of TM7 (**Fig. 7B**). This change in helical tilt occurred for the entire heptahelical bundle, with an increase in tilt for TM1, TM2, TM3, TM5, and TM7, and a decrease in tilt for TM4 and TM6 (**Fig. S3**). The longer C-terminus in the full-length A_2A_R permits greater rearrangements in the transmembrane regions, leading to the observed change in helical tilt. This result hints at potential conformational changes of A_2A_R upon oligomerization, necessitating future investigation on functional consequences.

C-terminal truncations prior to crystallization and structural studies may be the main reason for the scarcity of GPCR structures featuring oligomers. In that context, this study offers valuable insights and approaches to tune the oligomerization of A_2A_R and potentially of other GPCRs using its intrinsically disordered C-terminus. The presence of A_2A_R oligomeric populations with partial C-terminal truncations means that one can now study its oligomerization with less perturbation from the C-terminus. We also present evidence that the multiple C-terminal interactions that drive A_2A_R oligomerization can be easily modulated by ionic strength and specific salts (**Fig. 5** and **6**). Given that ~75% and ~15% of all class-A GPCRs possess a C-terminus of > 50 and > 100 amino acid residues(108), respectively, it will be worthwhile to explore the prospect of tuning GPCR oligomerization not only by shortening the C-terminus but also with simpler approaches such as modulating ionic strength and the surrounding salt environment.

## CONCLUSION

This study emphasizes for the first time the definite impact of the C-terminus on A_2A_R oligomerization, which can be extended to include the oligomers formed by other GPCRs with a protracted C-terminus. We have shown that the oligomerization of A_2A_R is strongly driven by depletion interactions along the C-terminus, further modulating and enhancing the multiple interfaces formed via a combination of hydrogen, electrostatic, hydrophobic, and covalent disulfide interactions. The task remains to link A_2A_R oligomerization to functional roles of the receptor(109). From a structural biology standpoint, visualizing the multiple oligomeric interfaces of A_2A_R in the presence of the full-length C-terminus is key to investigating whether these interfaces give rise to different oligomer functions.

## MATERIALS AND METHODS

### Cloning, Gene Expression, and Protein Purification

The multi-integrating pITy plasmid(110), previously used for overexpression of A_2A_R in *Saccharomyces cerevisiae*(111), was employed in this study. pITy contains a Gal1–10 promoter for galactose-induced expression, a synthetic pre-pro leader sequence which directs protein trafficking(112, 113), and the yeast alpha terminator. The genes encoding A_2A_R variants with 10-His C-terminal tag were cloned into pITy downstream of the pre-pro leader sequence, using either splice overlapping extension(114) or USER cloning using X7 polymerase(115, 116), with primers provided in **Table S3**. The plasmids were then transformed into *S. cerevisiae* strain BJ5464 (MATα ura3-52 trp1 leu2Δ1 his3Δ200 pep4::HIS3 prb1Δ1.6R can1 GAL) (provided by the lab of Anne Robinson at Carnegie Mellon University) using the lithium-acetate/PEG method(117). Transformants were selected on YPD G-418 plates (1% yeast extract, 2% peptone, 2% dextrose, 2.0 mg/mL G-418).

Receptor was expressed and purified following the previously described protocol(118). In brief, from freshly streaked YPD plates (1% yeast extract, 2% peptone, 2% dextrose), single colonies were grown in 5-mL YPD cultures over night at 30°C. From these 5-mL cultures, 50-mL cultures were grown with a starting OD of 0.5 over night at 30°C. To induce expression, yeast cells from these 50-mL cultures were centrifuged at 3,000 *x g* to remove YPD before resuspended in YPG medium (1% yeast, 2% peptone, 2% D-galactose) at a starting OD of 0.5. The receptor was expressed for 24 hours over night at 30°C with 250 r.p.m shaking. Cells were pelleted by centrifugation at 3,000 *x g*, washed in sterile PBS buffer, and pelleted again before storage at – 80°C until purification.

Mechanical bead lysis of cells was done, per 250 mL of cell culture, by performing 12 pulses of 60 s intense vortexing (with at least 60 s of rest in between pulses) in 10 mL 0.5-mm zirconia silica beads (BioSpec, Bartlesville, OK, USA; #11079105z), 25 mL of lysis buffer (50 mM sodium phosphate, 300 mM sodium chloride, 10% (v/v) glycerol, pH = 8.0, 2% (w/v) n-Dodecyl-β-D-maltopyranoside (DDM; Anatrace, Maumee, OH, USA; #D310), 1% (w/v) 3-[(3-Cholamidopropyl)dimethylammonio]-1-propanesulfonate (CHAPS; Anatrace; #C216), and 0.2% (w/v) cholesteryl hemisuccinate (CHS; Anatrace; #CH210) and an appropriate amount of 100x Pierce Halt EDTA-free protease inhibitor (Pierce, Rockford, IL, USA #78439)). Beads were separated using a Kontex column. Unlysed cells were removed by centrifugation at 3,220 *x g* for 10 min. Receptor was let solubilized on rotary mixer for 3 hours before cell debris was removed by centrifugation at 10,000 *x g* for 30 min. Solubilized protein was incubated with Ni-NTA resin (Pierce; #88221) over night. Protein-resin mixture was then washed extensively in purification buffer (50 mM sodium phosphate, 300 mM sodium chloride, 10% (v/v) glycerol, 0.1% (w/v) DDM, 0.1% (w/v) CHAPS and 0.2% (w/v) CHS, pH = 8.0) containing low imidazole concentrations (20– 50 mM). A_2A_R was eluted into purification buffer containing 500 mM imidazole. Prior to further chromatographic purification, imidazole was removed using a PD-10 desalting column (GE Healthcare, Pittsburgh, PA, USA; # 17085101).

Ligand affinity resin was prepared as previously described for purification of active A_2A_R.(119) (120) In brief, 8 mL of isopropanol-washed Affigel 10 resin (BioRad; # 1536099) was mixed gently in an Erlenmeyer flask for 20 h at room temperature with 48 mL of DMSO containing 24 mg of xanthine amine congener (XAC, high-affinity A_2A_R antagonist, K_D_ = 32 nM; Sigma, St. Louis, MO, USA; #X103). The absorbance at 310 nm of the XAC-DMSO solution before and after the coupling reaction was measured in 10 mM HCl and compared to a standard curve. The amount of resin bound to ligand was estimated to be 5.6 *μ*M. The coupling reaction was quenched by washing the resin with DMSO, then with Tris-HCl 50 mM (pH = 7.4), then with 20% (v/v) ethanol. The resin was packed into a Tricorn 10/50 column (GE Healthcare) under pressure via a BioRad Duoflow FPLC (BioRad).

For purification of active A_2A_R, the column was equilibrated with 4 CV of purification buffer. The IMAC-purified A_2A_R was desalted and diluted to 5.5 mL before applied to a 5-mL sample loop on the BioRad Duoflow FPLC, from which the sample was loaded onto the column at a rate of 0.1 mL/min. Inactive A_2A_R was washed from the column by flowing 10 mL of purification buffer at 0.2 mL/min, followed by 16 mL at 0.4 mL/min. Active A_2A_R was eluted from the column by flowing purification buffer containing 20 mM theophylline (low-affinity A_2A_R antagonist, K_D_ = 1.6 *μ*M; Sigma, St. Louis, MO, USA; #T1633). Western blot analysis was performed to determine 4-mL fractions with active A_2A_R collected with a BioFrac fraction collector (BioRad; Hercules, CA, USA), which were then concentrated through a 30-kDa MWCO centrifugal filter (Millipore, Billerica, MA, USA; # UFC803096) and desalted to remove excess theophylline. For the experiments where the salt concentrations were varied, the buffer exchange was done also by this last desalting step.

### Size-Exclusion Chromatography

To separate oligomeric species of active A_2A_R, a prepacked Tricorn Superdex 200 10/300 GL column (GE Healthcare) connected to a BioRad Duoflow FPLC was equilibrated with 60 mL of running buffer (150 mM sodium chloride, 50 mM sodium phosphate, 10% (v/v) glycerol, 0.1% (w/v) DDM, 0.1% (w/v) CHAPS, 0.02% (w/v) CHS, pH = 8.0) at a flow rate of 0.2 mL/min. 0.5-mL fractions were collected with a BioFrac fraction collector in 30 mL of running buffer at the same flow rate. Analysis of SDS/PAGE and western blot was done to determine oligomeric states of the eluted A_2A_R.

### SEC Peak Analysis

SEC chromatograms were analyzed using OriginLab using the nonlinear curve fit (Gaussian) function. The area under the curve and the peak width were manually defined in cases where the SNR of the SEC trace were too low. The R^2^ values reached > 0.96 for most cases. The population of each oligomeric species was expressed as the integral of each Gaussian this curve fit of the SEC signal. The HMW oligomer peak in some cases could not be fitted with one curve and thus was fitted with two curves instead. The reported standard errors were calculated from the variance of the fit and did not correspond to experimental errors. The results are detailed in **Fig. S2** and **Table S1**.

### SDS-PAGE and Western Blotting

10% SDS-PAGE gels were hand-casted in BioRad Criterion empty cassettes (BioRad; #3459902, 3459903). Lysate controls were prepared by lysis of 5 OD cell pellets with 35 *μ*L of YPER (Fisher Scientific, Waltham, MA, USA # 8990) at RT for 20 min, incubation with 2x Laemmli buffer (4% (w/v) SDS, 16% (v/v) glycerol, 0.02% (w/v) bromophenol blue, 167 M Tris, pH 6.8) at 37°C for 1 h, and centrifugation at 3,000 *x g* for 1 min to pellet cell debris. Protein samples were prepared by incubation with 2x Laemmli buffer at 37°C for 30 min. For all samples, 14 *μ*L (for 26-well gel) or 20 *μ*L (for 18-well gel) was loaded per lane, except for 7 *μ*L of Magic Mark XP Western protein ladder (Thermo Scientific, Waltham, MA, USA; # LC5602) as a standard. Electrophoresis was carried out at 120 V for 100 min. Proteins were transferred to 0.2-*μ*m nitrocellulose membranes (BioRad; # 170-4159) via electroblotting using a BioRad Transblot Turbo, mixed MW protocol. Membranes were blocked in Tris-buffered saline with Tween (TBST; 150 mM sodium chloride, 15.2 mM Tris-HCl, 4.6 mM Tris base, pH = 7.4, 0.1% (v/v) Tween 20 (BioRad; # 1706531)) containing 5% (w/v) dry milk, then probed with anti-A_2A_R antibody, clone 7F6-G5-A2 (Millipore, Burlington, MA, USA; # 05-717) at 1:500 in TBST with 0.5% (w/v) dry milk. Probing with secondary antibody was done with a fluorescent DyLight 550 antibody (Abcam, Cambridge, MA, USA; ab96880) at 1:600 in TBST containing 0.5% (w/v) milk.

Western blot was analyzed with Fiji. The Gels analysis plugin was used to define each sample lane, and to generate an intensity profile. Peaks were manually selected and integrated with the measure tool to determine the amount of protein present.

### Coarse-Grained MD Simulations

Initial configuration of A_2A_R was based on the crystal structure of the receptor in the active state (PDB 5G53). All non-receptor components were removed, and missing residues added using MODELLER 9.23(121). Default protonation states of ionizable residues were used. The resulting structure was converted to MARTINI coarse-grained topology using the martinize.py script(122). The ELNeDyn elastic network(123) was used to constrain protein secondary and tertiary structures with a force constant of 500 kJ/mol/nm^2^ and a cutoff of 1.5 nm. To optimize loop refinement of the model, a single copy was embedded in a 1-palmitoyl-2-oleoyl-sn-glycero-3-phosphocholine (POPC) bilayer using the insane.py script, solvated with MARTINI polarizable water, neutralized with 0.15 M NaCl, and a short MD (1.5 μs) run to equilibrate the loop regions. Subsequently, two monomers of the equilibrated A_2A_R were randomly rotated and placed at the center of a 13 nm × 13 nm × 11 nm (xyz) box, 3.5 nm apart, with their principal transmembrane axis aligned parallel to the z axis. The proteins were then embedded in a POPC bilayer using the insane.py script. Sodium and chloride ions were added to neutralize the system and obtain a concentration of 0.15 M NaCl. Total system size was typically in the range of 34,000 CG particles, with a 280:1 lipid:protein ratio. Ten independent copies were generated for each A_2A_R truncated variant.

v2.2 of the MARTINI coarse-grained force field(124) was used for the protein and water, and v2.0 was used for POPC. All coarse-grained simulations were carried out in GROMACS 2016(125) in the NPT ensemble (P = 1 atm, T = 310 K). The Bussi velocity rescaling thermostat was used for temperature control with a coupling constant of τ_t_ = 1.0 ps(126), while the Parrinello-Rahman barostat(127) was used to control the pressure semi-isotropically with a coupling constant of τ_t_ = 12.0 ps and compressibility of 3 × 10–4 bar–1. Reaction field electrostatics was used with Coulomb cut-off of 1.1 nm. Non-bonded Lennard-Jones interactions were treated with a cut-off of 1.1 nm. All simulations were run with a 15 fs timestep, updating neighbor lists every 10 steps. Cubic periodic boundary conditions along the x, y and z axes were used. Each simulation was run for 8 μs.

### Atomistic MD Simulations

Three snapshots of symmetric dimers of A_2A_R for each respective truncated variant were randomly selected from the CG simulations as starting structures for backmapping. Coarse-grained systems were converted to atomistic resolution using the backward.py script(128). All simulations were run in Gromacs2019 in the *NPT* ensemble (*P* = 1 bar, *T* = 310 K) with all bonds restrained using the LINCS method(129). The Parrinello-Rahman barostat was used to control the pressure semi-isotropically with a coupling constant of τ_t_ = 1.0 ps and a compressibility of 4.5 × 10–5 bar–1, while the Bussi velocity rescaling thermostat was used for temperature control with a coupling constant of τ_t_ = 0.1 ps. Proteins, lipids, and solvents were separately coupled to the thermostat. The CHARMM36 and TIP3P force fields(130, 131) were used to model all molecular interactions. Periodic boundary conditions were set in the x, y, and z directions. Particle mesh Ewald (PME) electrostatics was used with a cut-off of 1.0 nm. A 2-fs time step was used for all atomistic runs, and each simulation was run for 50 ns.

### Analysis of Computational Results

All trajectories were post-processed using gromacs tools and in-house scripts. We ran a clustering analysis of all dimer frames from the CG simulations using Daura et. al.’s clustering algorithm(132) implemented in GROMACS, with an RMSD cutoff of 1.5 Å. (An interface was considered dimeric if the minimum center of mass distance between the protomers was less than 5 Å.) This method uses an RMSD cutoff to group all conformations with the largest number of neighbors into a cluster and eliminates these from the pool, then repeats the process until the pool is empty. We focused our analysis on the most populated cluster from each truncated variant. Electrostatic interactions in the dimer were calculated from CG systems with LOOS(133) using a distance cutoff of 5.0 Å.

Transmembrane helical tilt angles were also calculated in LOOS from CG simulations. Hydrogen bonds were calculated from AA simulations using the hydrogen bonds plugin in VMD(134), with a distance cutoff of 3.5 Å and an angle cutoff of 20°. Only C-terminal residues were included in hydrogen bond analysis. PyMOL(135) was used for molecular visualizations.

### Assessing A_2A_R Oligomerization with Increasing Ionic Strength

Na_2_HPO_4_ and NaH_2_PO_4_ in the buffer make up an ionic strength of 0.15 M, to which NaCl was added to increase the ionic strength to 0.45 M and furthermore to 0.95 M. The A_2A_R variants were purified at 0.45 M ionic strength and then exchanged into buffers of different ionic strengths using a PD-10 desalting column prior to subjecting the samples to SEC. The buffer composition is detailed below.

### Isolated C-Terminus Purification

*Escherichia coli* BL21 (DE3) cells were transfected with pET28a DNA plasmids containing the desired A_2A_R sequence with a 6x His tag attached for purification. Cells from glycerol stock were grown in 10 mL luria broth (LB, Sigma Aldrich, L3022) overnight at 37°C and then used to inoculate 1 L of fresh LB and 10 μg/mL kanamycin (Fisher Scientific, BP906). Growth of cells were performed at 37°C, 200 rpm until optical density at λ = 600 nm reached 0.6–0.8. Expression was induced by incubation with 1 mM isopropyl-β-D-thiogalactoside (Fisher Bioreagents, BP175510) for 3 hrs.

Cells were harvested with centrifugation at 5000 rpm for 30 min. Harvested cells were resuspended in 25 mL Tris-HCl, pH = 7.4, 100 mM NaCl, 0.5 mM DTT, 0.1 mM EDTA with 1 Pierce protease inhibitor tablet (Thermo Scientific, A32965), 1 mM PMSF, 2 mg/mL lysozyme, 20 *μ*g/mL DNase (Sigma, DN25) and 10 mM MgCl2, and incubated on ice for 30 min. Samples were then incubated at 30°C for 20 minutes, then flash frozen and thawed 3 times in LN2. Samples were then centrifuged at 10,000 rpm for 10 min to remove cell debris. 1 mM PMSF was added again and the resulting supernatant was incubated while rotating for at least 4 hrs with Ni-NTA resin. The resin was loaded to a column and washed with 25 mL 20 mM sodium phosphate, pH = 7.0, 1 M NaCl, 20 mM imidazole, 0.5 mM DTT, 100 *μ*M EDTA. Purified protein was eluted with 15 mL of 20 mM sodium phosphate, pH = 7.0, 0.5 mM DTT, 100 mM NaCl, 300 mM imidazole. The protein was concentrated to a volume of 2.5mL and was buffer exchanged into 20 mM ammonium acetate buffer, pH = 7.4, 100 mM NaCl using a GE PD-10 desalting column. Purity of sample was confirmed with SDS-PAGE and western blot.

### Aggregation Assay to Assess A_2A_R C-Terminus Assembly

Absorbance was measured at 450 nm using a Shimadzu UV-1601 spectrophotometer with 120 *μ*L sample size. Prior to reading, samples were incubated at 40°C for 5 minutes. Samples were vigorously pipetted to homogenize any precipitate before absorbance was measured. Protein concentration was 50 *μ*M in a 20 mM ammonium acetate buffer (pH = 7.4).

### Differential Scanning Fluorimetry (DSF)

DSF was conducted with a Bio-rad CFX90 real-time PCR machine. A starting temperature 20°C was increased at a rate of 0.5°C per 30 seconds to a final temperature of 85°C. All samples contained 40 μL of 40 *μ*M A_2A_R C-terminus, 9x SYPRO orange (ThermoFisher S6650), 200 mM NaCl, and 20 mM MES. Fluorescence was detected in real-time at 570 nm. All samples were conducted in triplicate.

### Hydrophobicity and Charge Profile of C-Terminus

The hydrophobicity profile reported in **Fig. S4** was determined with ProtScale using method described by Kyte & Doolittle(136), window size of 3.

## FUNDING AND ACKNOWLEDGMENTS

This material is based upon work supported by (1) the National Institute of General Medical Sciences of the National Institutes of Health under Award Number R35GM136411, (2) the National Institute of Mental Health of the National Institutes of Health under Small Business Innovation Research Award Number 1R43MH119906-01, and (3) the National Science Foundation under Award Number MCB-1714888 (E.S. and B.M.). The content is solely the responsibility of the authors and does not necessarily represent the official views of the National Institutes of Health. Many of the experiments were completed with the assistance from Rohan Katpally. The pITy expression vector and *S. cerevisiae* BJ5464 strain were generously provided by Prof. Anne Robinson’s lab at Carnegie Mellon University. The X7 polymerase was a gift from Dr. Morten Nørholm, Novo Nordisk Foundation Center for Biosustainability, Technical University of Denmark. Computational time was provided through WVU Research Computing and XSEDE allocation no. TG-MCB130040.

## Notes

### Competing Interest Statement

The authors have declared no competing interest.

