## Supporting Information for "Oligomerization of the Human Adenosine A_2A_ Receptor is Driven by the Intrinsically Disordered C-terminus"

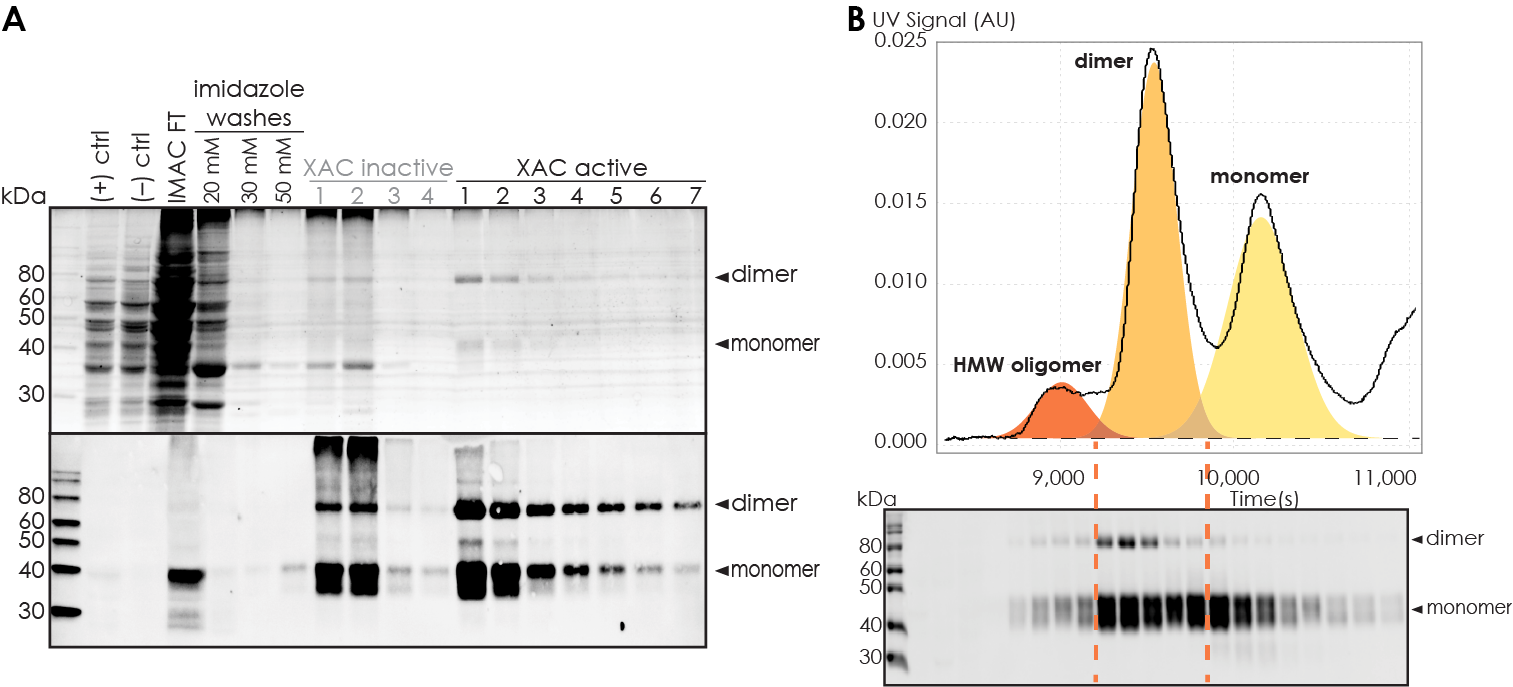

**Figure S1. (A)** Representative total protein stain (upper panel) and western blot (lower panel) of A_2A_R-WT during purification. Positive ((+) ctrl) and negative ((–) ctrl) controls consist of 5 OD cell lysate of S. cerevisiae BJ5464 cells expressing and not expressing A_2A_R WT, respectively. “IMAC FT” indicates the flow-through from IMAC step. “XAC inactive” and “XAC active” indicate the fractions that do not and do bind to XAC during the ligand-affinity chromatography step. **(B)** Representative western blot of A_2A_R-WT during SEC separation. The fractions are matched to the distinct oligomeric peaks in the SEC chromatogram. Each lane on the blot is from 0.5 mL fractions eluted from a Superdex 200 10/300 GL (GE Healthcare) column. MagicMark protein ladder (LC5602) is used as the molecular weight standard.

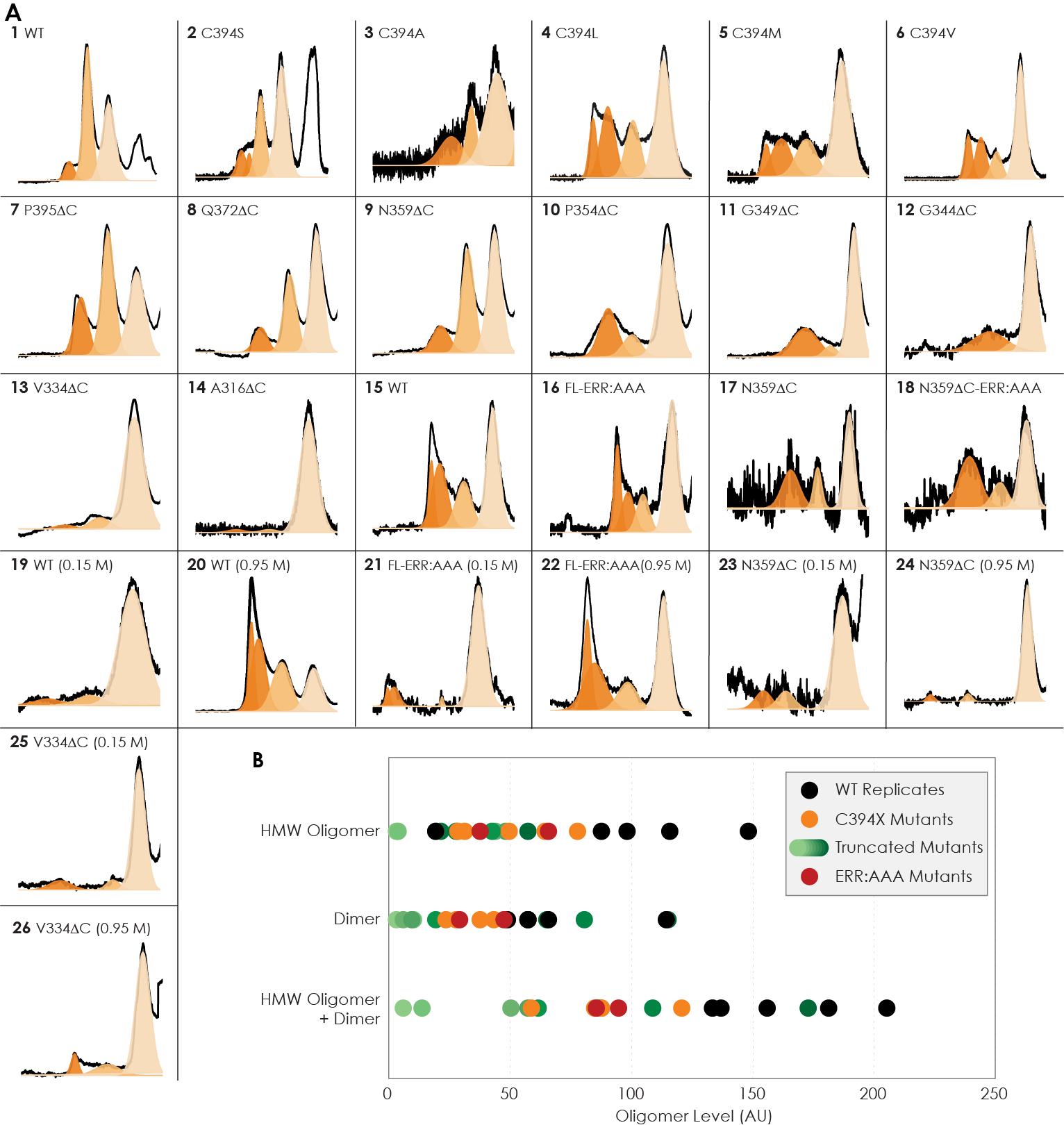

**Figure S2. (A)** Curve fitting using OriginLab of all A_2A_R variants used in this study. By default, each oligomeric peak is fitted with one curve using Gaussian distribution and displayed by different color shades, with the HMW oligomer eluted first (dark orange), followed by the dimer (lighter orange), followed by the monomer (lightest orange). However, the HMW oligomer peak in some cases cannot be fitted with one curve and thus is fitted with two curves instead. This discrepancy can be explained by variation in HMW oligomerization order among the variants. The identity of each peak is confirmed with western blotting. The value and error from the curve fitting of each peak are given in **Table S1**. **(B)** Data distribution of all variants used in this study in comparison to five experimental replicates of A_2A_R-WT. The C-terminally truncated mutants are represented by different shades of green in increasing darkness corresponding to the increased length of the C-terminus, with the lightest shade representing the mutant with the shortest C-terminus (A316ΔC) and the darkest shade for the mutant with the longest C-terminus (P395ΔC). The levels of dimer and HMW oligomer are expressed relative to the monomeric population in arbitrary unit, with reported errors calculated from the variance of the fit, not experimental variation. There are significant variations in the dimer and HMW oligomer levels among the WT replicates, stemming from experimental errors. These variations are mitigated when the two parameters are added, as the data distribution becomes more uniform. Also, the oligomerization levels of the WT replicates are consistently higher than the mutated and truncated variants.

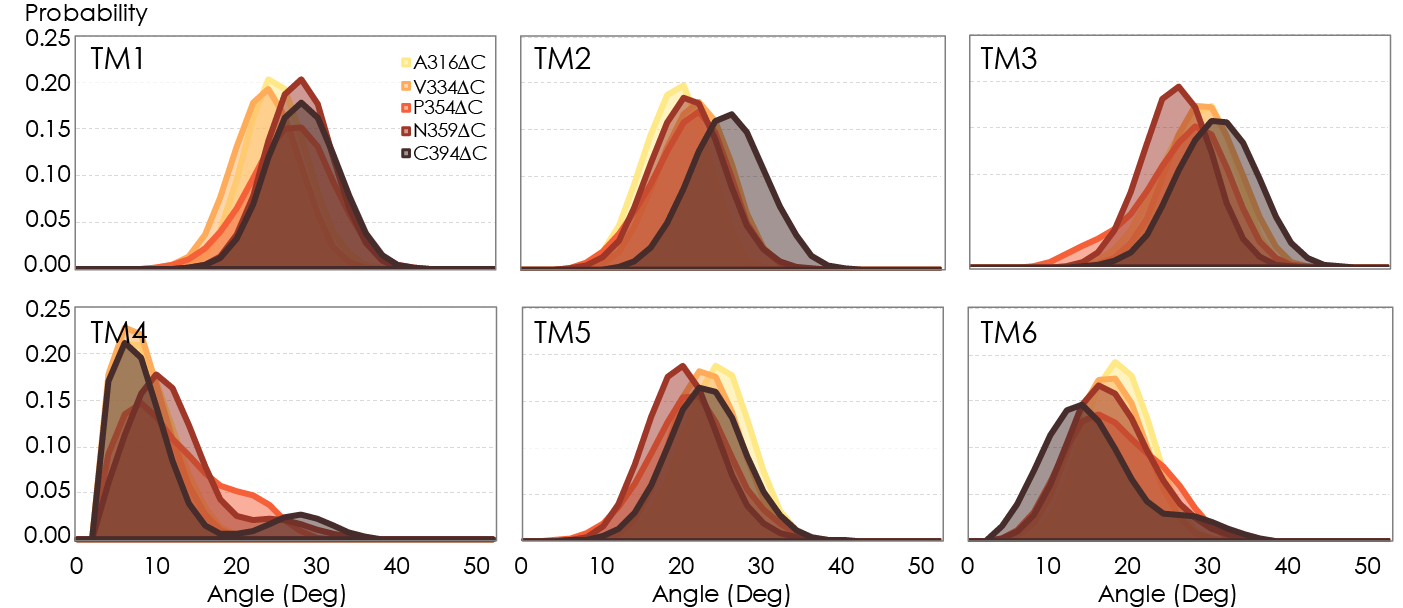

**Figure S3.** Helical tilt angles for TM1­–6 helices in A_2A_R as a function of protein length. Systematic truncations of the C-terminus lead to rearrangement of the heptahelical bundle, propagated to the entire receptor and is especially pronounced in helices proximal to the C-terminus, i.e. TM1, TM2, TM7. For almost all TM helices, a noticeable shift in tilt angle occurs upon modeling the full-length (394 residues) variant. This behavior is fundamentally different from the conventional model of GPCR activation, in which TM 1, 2, 4, and 7 remain fairly rigid, with TM5 and TM6 undergoing an outward tilt/rotation to enable binding to the cognate G protein. Relaxation of the heptahelical bundle (i.e., an increase in helical tilt) as a function of protein length and dimerization could potentially be critical to our understanding of the activation mechanism of A_2A_R, as past studies have overwhelmingly focused on activation of the monomer.

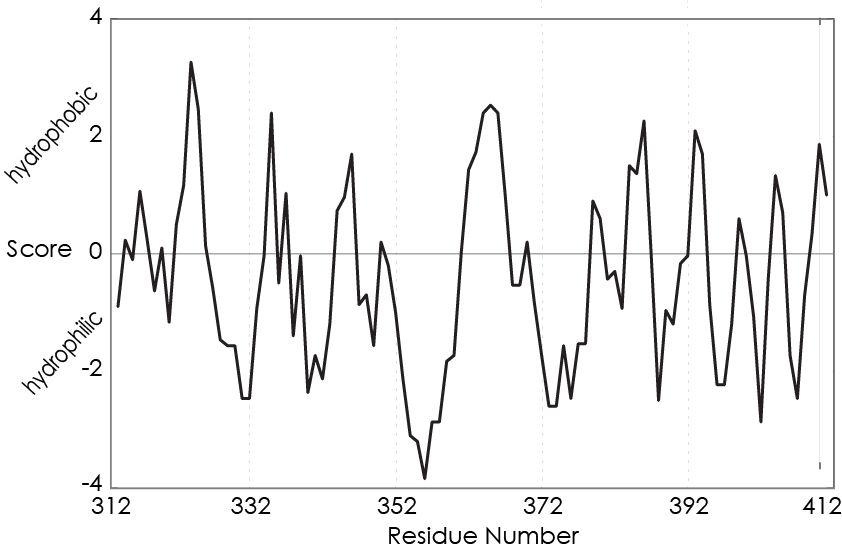

**Figure S4.** Hydropathy plot against A_2A_R residue number showing the hydrophobicity of A_2A_R C-terminus, scored with ProtScale using method described by Kyte & Doolittle, window size of 3. Positive scores represent hydrophobicity and negative scores hydrophilicity.

| **Fig** | **Variants** | **No.** | **HMW Oligomer Level** | **Dimer Level** | **Total Oligomer Level** | **[HMW Oligomer]** | **[Dimer]** | **[Monomer]** |
| --- | --- | --- | --- | --- | --- | --- | --- | --- |
| 2 | WT | 1 | 0.20 ± 0.01 | 1.14 ± 0.01 | 1.34 ± 0.01 | 1.77 ± 0.05 | 10.39 ± 0.05 | 9.09 ± 0.07 |
|  | C394S | 2 | 0.28 ± 0.06 | 0.57 ± 0.01 | 0.85 ± 0.06 | 1.66 ± 0.35 | 3.36 ± 0.07 | 5.90 ± 0.06 |
|  | C394A | 3 | 0.31 ± 0.08 | 0.28 ± 0.06 | 0.59 ± 0.10 | 0.49 ± 0.11 | 0.44 ± 0.10 | 1.57 ± 0.08 |
|  | C394L | 4 | 0.78 ± 0.01 | 0.43 ± 0.01 | 1.21 ± 0.01 | 9.09 ± 0.13 | 5.07 ± 0.07 | 11.73 ± 0.09 |
|  | C394M | 5 | 0.50 ± 0.08 | 0.38 ± 0.03 | 0.88 ± 0.09 | 2.70 ± 0.42 | 2.05 ± 0.18 | 5.44 ± 0.05 |
|  | C394V | 6 | 0.64 ± 0.01 | 0.23 ± 0.01 | 0.88 ± 0.01 | 9.94 ± 0.13 | 3.65 ± 0.06 | 15.44 ± 0.07 |
| 3 | WT | 1 | 0.20 ± 0.01 | 1.14 ± 0.01 | 1.34 ± 0.01 | 1.77 ± 0.05 | 10.39 ± 0.05 | 9.09 ± 0.07 |
|  | P395ΔC | 7 | 0.58 ± 0.01 | 1.15 ± 0.01 | 1.73 ± 0.02 | 3.34 ± 0.05 | 6.69 ± 0.05 | 5.80 ± 0.06 |
|  | Q372ΔC | 8 | 0.22 ± 0.01 | 0.65 ± 0.01 | 0.87 ± 0.01 | 1.64 ± 0.05 | 4.95 ± 0.05 | 7.59 ± 0.06 |
|  | N359ΔC | 9 | 0.28 ± 0.01 | 0.81 ± 0.01 | 1.09 ± 0.01 | 2.31 ± 0.06 | 6.72 ± 0.05 | 8.30 ± 0.06 |
|  | P354ΔC | 10 | 0.42 ± 0.01 | 0.19 ± 0.01 | 0.62 ± 0.02 | 2.17 ± 0.05 | 0.99 ± 0.05 | 5.12 ± 0.05 |
|  | G349ΔC | 11 | 0.48 ± 0.02 | 0.09 ± 0.01 | 0.58 ± 0.02 | 2.23 ± 0.07 | 0.42 ± 0.06 | 4.60 ± 0.03 |
|  | G344ΔC | 12 | 0.44 ± 0.10 | 0.06 ± 0.06 | 0.50 ± 0.12 | 0.80 ± 0.18 | 0.11 ± 0.11 | 1.81 ± 0.04 |
|  | V334ΔC | 13 | 0.04 ± 0.01 | 0.10 ± 0.01 | 0.14 ± 0.01 | 0.29 ± 0.06 | 0.83 ± 0.06 | 8.23 ± 0.06 |
|  | A316ΔC | 14 | 0.03 ± 0.01 | 0.03 ± 0.01 | 0.06 ± 0.01 | 0.08 ± 0.02 | 0.08 ± 0.02 | 2.89 ± 0.02 |
| 4 | WT | 15 | 0.88 ± 0.04 | 0.49 ± 0.01 | 1.37 ± 0.01 | 5.37 ± 0.22 | 2.98 ± 0.07 | 6.10 ± 0.04 |
|  | WT-ERRAAA | 16 | 0.66 ± 0.03 | 0.29 ± 0.01 | 0.95 ± 0.03 | 3.76 ± 0.16 | 1.64 ± 0.08 | 5.72 ± 0.07 |
|  | N359ΔC | 17 | 0.68 ± 0.04 | 0.33 ± 0.03 | 1.01 ± 0.05 | 1.10 ± 0.06 | 0.53 ± 0.04 | 1.61 ± 0.04 |
|  | N359ΔC-ERRAAA | 18 | 0.38 ± 0.03 | 0.48 ± 0.02 | 0.85 ± 0.04 | 1.05 ± 0.08 | 1.32 ± 0.06 | 2.78 ± 0.05 |
| 5 | WT 0.15 M | 19 | 0.07 ± 0.01 | 0.09 ± 0.01 | 0.16 ± 0.02 | 0.19 ± 0.04 | 0.27 ± 0.04 | 2.87 ± 0.04 |
|  | WT 0.45 M | 15 | 0.88 ± 0.04 | 0.49 ± 0.01 | 1.37 ± 0.04 | 5.37 ± 0.22 | 2.98 ± 0.07 | 6.10 ± 0.04 |
|  | WT 0.95 M | 20 | 2.20 ± 0.04 | 1.31 ± 0.02 | 3.51 ± 0.05 | 14.54 ± 0.25 | 8.62 ± 0.11 | 6.60 ± 0.06 |
|  | WT-ERRAAA 0.15 M | 21 | 0.17 ± 0.05 | 0.02 ± 0.01 | 0.19 ± 0.05 | 0.62 ± 0.17 | 0.07 ± 0.01 | 3.73 ± 0.03 |
|  | WT-ERRAAA 0.45 M | 16 | 0.47 ± 0.08 | 0.45 ± 0.04 | 0.92 ± 0.09 | 2.55 ± 0.45 | 2.45 ± 0.23 | 5.45 ± 0.07 |
|  | WT-ERRAAA 0.95 M | 22 | 1.20 ± 0.03 | 0.38 ± 0.01 | 1.58 ± 0.03 | 7.41 ± 0.18 | 2.37 ± 0.08 | 6.21 ± 0.04 |
|  | N359ΔC 0.15 M | 23 | 0.11 ± 0.01 | 0.11 ± 0.01 | 0.21 ± 0.02 | 0.72 ± 0.08 | 0.71 ± 0.08 | 6.67 ± 0.07 |
|  | N359ΔC 0.45 M | 17 | 0.68 ± 0.04 | 0.33 ± 0.03 | 1.01 ± 0.05 | 1.10 ± 0.06 | 0.53 ± 0.04 | 1.61 ± 0.04 |
|  | N359ΔC 0.95 M | 24 | 0.04 ± 0.01 | 0.04 ± 0.01 | 0.09 ± 0.01 | 0.51 ± 0.05 | 0.59 ± 0.05 | 11.90 ± 0.06 |
|  | V334ΔC 0.15 M | 25 | 0.13 ± 0.01 | 0.08 ± 0.01 | 0.21 ± 0.01 | 0.65 ± 0.04 | 0.41 ± 0.03 | 5.03 ± 0.03 |
|  | V334ΔC 0.45 M | 8 | 0.04 ± 0.01 | 0.10 ± 0.01 | 0.14 ± 0.01 | 0.29 ± 0.06 | 0.83 ± 0.06 | 8.23 ± 0.06 |
|  | V334ΔC 0.95 M | 26 | 0.09 ± 0.02 | 0.15 ± 0.04 | 0.23 ± 0.01 | 0.85 ± 0.19 | 1.41 ± 0.34 | 9.68 ± 0.27 |
| WT Replicates | | | 1.16 ± 0.05 | 0.65 ± 0.03 | 1.81 ± 0.06 | 9.45 ± 0.39 | 5.34 ± 0.20 | 8.16 ± 0.04 |
|  |  |  | 0.98 ± 0.03 | 0.57 ± 0.01 | 1.56 ± 0.04 | 6.44 ± 0.20 | 3.76 ± 0.09 | 6.55 ± 0.04 |
|  |  |  | 1.48 ± 0.05 | 0.57 ± 0.01 | 2.05 ± 0.05 | 12.02 ± 0.35 | 4.66 ± 0.06 | 8.12 ± 0.05 |
|  |  |  | 0.20 ± 0.01 | 1.14 ± 0.01 | 1.34 ± 0.01 | 1.77 ± 0.05 | 10.39 ± 0.05 | 9.09 ± 0.07 |
|  |  |  | 0.88 ± 0.04 | 0.49 ± 0.01 | 1.37 ± 0.04 | 5.37 ± 0.22 | 2.98 ± 0.07 | 6.10 ± 0.04 |

**Table S1.** Results from curve fitting using OriginLab and calculations of the HMW oligomer and dimer levels for all A_2A_R variants used in this study. The variants are grouped by the order they appear in the main text and numbered corresponding to **Fig. S2**. The levels of dimer and HMW oligomer are expressed relative to the monomeric population in arbitrary units as monomer-equivalent concentration ratios. The errors are calculated from the variance of the fit, not experimental variations, and are within 95% confidence interval.
